# DNA 5-methylcytosine regulates genome-wide formation of G-quadruplex structures

**DOI:** 10.1101/2023.02.16.528796

**Authors:** Kangkang Niu, Lijun Xiang, Xiaoyu Li, Jin Li, Yuanli Li, Chu Zhang, Junpeng Liu, Xiaojuan Zhang, Yuling Peng, Guanfeng Xu, Hui Xiang, Hao Wang, Qisheng Song, Qili Feng

## Abstract

G-quadruplex structures (G4s) have been identified in genomes of multiple organisms and proven to play important epigenetic regulatory roles in various cellular functions. However, the G4 formation mechanism remains largely unknown. Here, we found a negative correlation between DNA 5mC methylation and G4 abundance. The abundance of genomic G4s significantly increased when the whole-genome methylation level was reduced in DNMT1-knockout cells. This increase was then suppressed by DNMT1 over-expression. And more G4s were detected in the hypomethylated cancer cell line HepG2 and rectal cancer tissues. Besides, 5mC modification significantly inhibited G4 formation of the potential G-quadruplex sequences (PQSs). The transcription of genes with 5mC modification sites in their promoter PQSs was affected after treatment with G4 stabilizer pyridostatin or methylation inhibitor 5-aza-dC. The global reduction of genomic methylation elevates gene transcription levels through increased G4s. Taken together, DNA 5mC methylation prevents PQSs from folding into G4s in genomes.

## INTRODUCTION

In addition to the typical DNA double-helix structure (Watson and Crick 1953), tetraplex DNA secondary structures have been reported in many studies (Choi and Majima 2011). DNA G-quadruplex (G4), a tetraplex structure, typically forms in guanine-rich sequences (Kwok and Merrick 2017; Raguseo et al. 2020; Spiegel et al. 2020). Four guanines are mutually connected by Hoogsteen hydrogen bonds to form a G-tetrad, which is stabilized by monovalent cations, such as K^+^, and two or more G-tetrads stack into a G4 structure (Sundquist and Klug 1989; Williamson et al. 1989; Dingley et al. 2005).

Based on the potential G-quadruplex sequence (PQS) characteristics (G_≥2_N_1-12_ G_≥2_N_1-12_G_≥2_N_1-12_G_≥2_N_1-12_; N: A, T or C), whole-genome analysis of PQS was performed for 37 species (Wu et al. 2021). A large number of PQSs were identified in the genomes of these species, and the number, frequency, density and length of G4s generally increased with genome evolution (Wu et al. 2021). Whole-genome mapping of G4s by G4 high-throughput sequencing method (G4-seq) was reported for 12 species, and 131 to 705,580 G4s were identified from *Escherichia coli* to *Homo sapiens* (Marsico et al. 2019). G4 structures were also visualized (Biffi et al. 2013) and identified (Hänsel-Hertsch et al. 2016, 2018) in human cells by using a BG4 antibody. A recent study reported an artificial G4 probe (G4P) protein that binds G4s with high affinity and specificity, and using this G4P probe, G4s were found in >60% of promoters and ~70% of genes in human cells (Zheng et al. 2020). Although elaborative cellular functions of G4s remain to be clarified, the involvement of these structures in different cellular functions has been reported (Rhodes and Lipps 2015; Tian et al. 2018; Varshney et al. 2020). G4s in the promoter of a gene can regulate transcription by recruiting transcription factors or blocking their access to the promoter (González et al. 2009; Borgognone et al. 2010; Li et al. 2017; Niu et al. 2019). G4s at telomeres can inhibit telomerase activity and lead to telomere shortening (Zahler et al. 1991; Read et al. 2001; Burger et al. 2005). Stabilization of G4 structures by G4 ligands or the absence of a G4-unwinding helicase can lead to genome instability by blocking DNA replication (Piazza et al. 2010, 2017). Collectively, G4s have been demonstrated to involve in many cellular functions in different species. One obvious and interesting question is what regulates the formation of G4s in genomes.

Methylation of the fifth position of cytosine (5mC) in DNA molecules is a widely studied epigenetic modification of DNA (Smith and Meissner 2013). In mammals, three DNA methyltransferases (DNMTs) have been reported to participate in DNA methylation. DNMT3A and DNMT3B catalyze *de novo* DNA methylation, while DNMT1 maintains methylation (Li et al. 1992; Jones 2012; Liao et al. 2015). Approximately 60-90% of CpGs in the entire human genome are methylated throughout lifetime (Lister et al. 2009), and CpG methylation is closely associated with transcriptional regulation, development and disease (Feinberg and Tycko 2004; Smith and Meissner 2013). DNA methylation generally can silence gene transcription by suppressing the binding of transcription factors to gene promoters (Klose and Bird 2006; Amabile et al. 2016; Liu et al. 2016; Vojta et al. 2016; Stepper et al. 2017). DNA methylation can also activate gene transcription by recruiting transcription factors that preferentially bind to methylated DNA (Yin et al. 2017). It seems that DNA methylation has a dual function in gene transcription regulation. Notably, dysfunction of DNA methylation may lead to human disease, especially cancer, probably by regulating the expression of specific oncogenes. Aberrant hypermethylation of tumor suppressor genes results in inactivation of these genes (Jones and Baylin 2002) and. Genome-wide aberrant hypomethylation can lead to a wide variety of tumors (Lapeyre and Becker 1979; Feinberg and Vogelstein 1983; Gama-Sosa et al. 1983; Ehrlich 2002; Gaudet et al. 2003).

An interesting issue is that many PQSs harbor in CpG islands (CGIs) and these sites are capable of either forming G4 structures or being methylated. This raises a question of whether an intrinsic interaction occurs between G4 formation and DNA methylation. In fact, a few studies have reported that G4 structures, especially those formed in CGIs, prevent the methylation of CpG dinucleotides (Halder et al. 2010; Mao et al. 2018). However, conversely, whether DNA methylation affects G4 formation remains unclear.

In this study, we used different approaches to determine the effects of DNA methylation on G4 formation at genome level and revealed a negative relationship between DNA methylation and G4 formation. Suppressing whole-genome methylation led to a genome-wide increase in the abundance of G4 structures.

## RESULTS

### More Abundance of G4s Formed in Hypomethylated Sequences

To investigate the effect of DNA methylation on the formation of G4 structures, methylation modification data and G4 CUT&Tag data from human HeLa and K562 cancer cells were obtained from the NCBI database, and the correlation between these two epigenetic mechanisms was analyzed.

Genes were sorted by methylation level, based on the data from whole-genome bisulfite sequencing (WGBS) of HeLa cells, in the 1 kb upstream and downstream regions around their transcription starting site (TSS). The top 10,000 hypermethylated genes (after removing duplicates, 6,002 genes remained) and the bottom 10,000 hypomethylated genes (after removing duplicates, 6,272 genes remained) were selected to perform the following analysis. Using the reported G4 CUT&Tag data for HeLa cells (Li et al. 2021), the G4 distribution in the same 1 kb upstream and downstream regions around the TSS of the selected genes was mapped. The results revealed that the G4 reads for the hypermethylated genes were significantly lower than those for all genes. The G4 reads for the hypomethylated genes were notably higher than those for all genes (Fig. 1A and B). To ensure that these differences were not caused by the intrinsic quantity variance of PQSs, we further analyzed whether the hypermethylated and hypomethylated modified genes contained similar amounts of PQSs. PQS mapping revealed that a similar amount of PQSs existed between those two gene sets, with slightly more PQSs in the hypermethylated genes (Fig. 1A and C).

**Figure 1.**
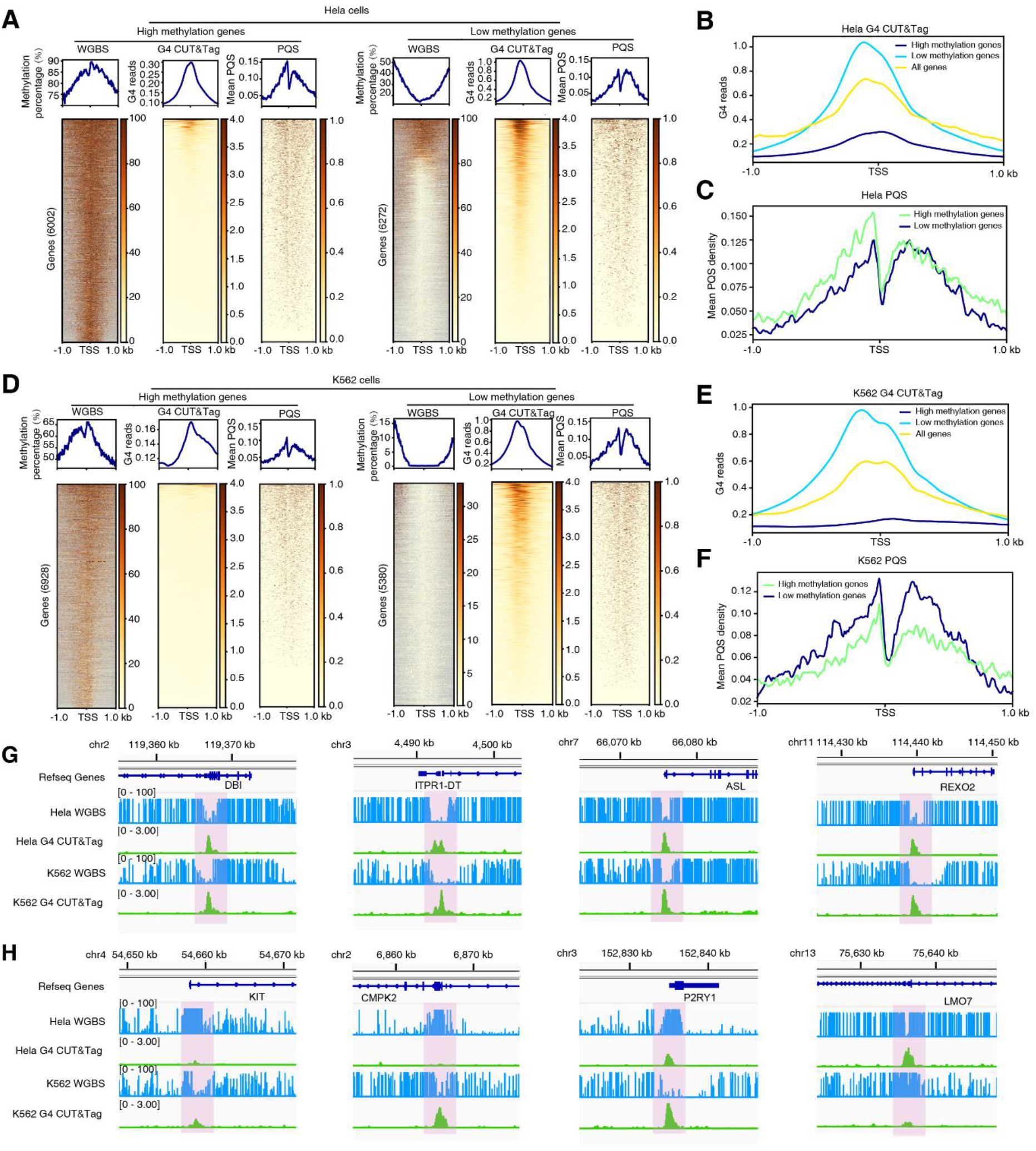
Comparison of DNA methylation and G4 formation in the 1 kb upstream and downstream regions around the TSSs of human genes. (A, D) Heatmap analysis of methylation, G4s and PQSs in HeLa and K562 cells. (B, E) The distribution of the G4 structures in the 1 kb upstream and downstream regions around the TSSs of hypermethylated and hypomethylated genes. (C, F) The PQS distribution of hypermethylated and hypomethylated genes. (G) IGV (Integrative Genomics Viewer) screen shots showing the coincidence of G4 peaks (green) with hypomethylated sites in HeLa and K562 cells. The whole genome bisulfite sequence (WGBS) tracks are in blue. (H) The IGV screen shots showing that in HeLa cells, the KIT, CMPK2 and P2RY1 sites had a high level of methylation and low G4 enrichment, while the LMO7 sites had a low level of methylation and high G4 enrichment. In K562 cells, the abundance of methylation modification of KIT, CMPK2 and P2RY1 decreased, and the enrichment of G4 increased, whereas the abundance of methylation modification of LMO7 increased, and the enrichment of G4 decreased.

A similar analysis was conducted with the data from K562 cells. Consistent with that in HeLa cells, the G4 read value was approximately ten times higher for the hypomethylated genes than that for the hypermethylated genes in K562 cells, although the PQS abundance in the hypomethylated genes was slightly higher than that in the hypermethylated genes (Fig. 1D-F). These results indicated that a negative relationship between G4 formation and DNA methylation exists in human genome. To validate this finding, eight fragments that contain PQSs were randomly selected from the two gene sets to perform further analysis. Chromosome alignment showed that in both HeLa and K562 cells, G4s were significantly enriched in hypomethylated regions (Fig. 1G). In HeLa cells, the KIT, CMPK2 and P2RY1 sites had a high level of methylation and a low G4 enrichment rate, while the LMO7 sites had a low level of methylation and a high G4 enrichment rate (Fig. 1H). However, in K562 cells, the methylation modification of KIT, CMPK2 and P2RY1 decreased, while the enrichment rate of G4 increased significantly. The methylation modification of LMO7 increased, and the enrichment rate of G4 decreased significantly (Fig. 1H).

These results imply that the PQSs within the hypermethylated regions cannot easily fold into G4 structures, but the PQSs in the hypomethylated regions can, suggesting that DNA methylation may block the formation of G4 structures.

### More Abundance of G4s Detected in Less Methylated Tumor Cells

Reduced genome-wide methylation is a typical feature of oncogenesis (Lapeyre and Becker 1979; Feinberg and Vogelstein 1983; Gama-Sosa et al. 1983; Ehrlich 2002; Gaudet et al. 2003). The above results indicated that hypomethylation may allow the formation of more G4 structures. To further verify this finding, methylation and G4 levels were measured in the nontumor cell line 293T derived from human embryonic kidney cells and the human hepatic adenocarcinoma cell line HepG2. The number of genomic G4s in HepG2 cells was significantly higher than that in 293T cells (Fig. 2A and B), while the overall methylation level in HepG2 cells was significantly lower than that in 293T cells (Fig. 2C). To exclude the effect of the cell type difference, a similar analysis was performed with cancerous and noncancerous rectal tissues. The results showed that the overall genomic methylation level of the cancerous rectal tissue was significantly lower than that of the noncancerous rectal tissue (Fig. 2E), and the abundance of genomic G4s in the cancerous tissues was significantly higher than that in the noncancerous tissues (Fig. 2D and F). All these evidence pointed out a negative correlation between DNA methylation and G4 formation at the genomic level.

**Figure 2.**
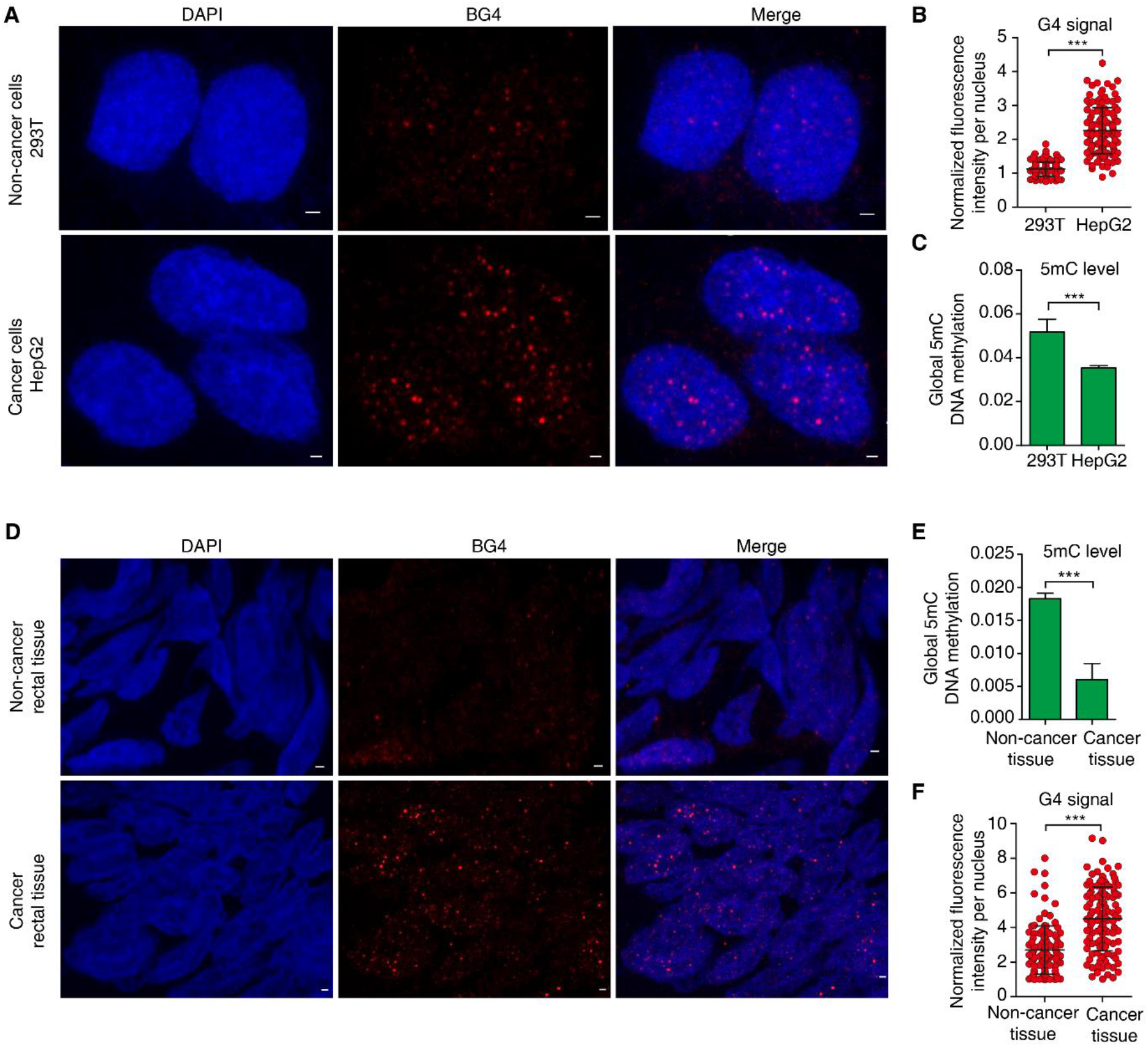
Comparative analysis of methylation and G4 levels between oncogenic and non-oncogenic cells or tissues. (A, D) The detection (A) and quantitative analysis (D) of G4 structures in 293T and HepG2 cells. (B, F) The detection (B) and quantitative analysis (F) of G4 structures in cancer rectal tissue and non-cancer rectal tissue. (C, E) The methylation level detected in 293T and HepG2 cells (C) and in cancerous rectal tissue and noncancerous rectal tissue (E). Data shown are the mean ± SD (D and F) or mean ± SEM (C and E). Statistical significance was determined by Student’s t test, ***p<0.001. The scale bars equal 1 μm.

### Abundance of G4s Significantly Increased When Methylation was Inhibited

DNA methylation is an epigenetic modification that is crucial for the development and differentiation of various cell types (Jones 2012; Smith and Meissner 2013). Inhibition of methylation has been implemented to clarify the critical role of DNA methylation. 5-Aza-dC and RG108 are well-known methylation inhibitors. 5-Aza-dC is a cytosine analog that inhibits DNMT functions by covalently binding to cysteine residues in DNA molecules (Scott et al. 2017). RG108 is a nonnucleoside DNMT inhibitor that blocks the DNMT active site (Brueckner et al. 2005).

To further investigate the effect of DNA methylation on G4 structure formation, the G4 signal variation in 293T and SK-Hep-1 cells was detected when the methylation level was inhibited with methylation inhibitors. In 293T cells, the methylation level was significantly decreased when the cells were treated with 5 μM 5-aza-dC (Fig. 3A and C), and correspondingly, the G4 signal was significantly increased (Fig. 3B and D). In addition, when human SK-Hep-1 cells were treated with RG108, the overall methylation level of the genome was significantly reduced (Fig. 3E and G), with a remarkable increase in the number of G4 structures (Fig. 3F and H). Thus, methylation inhibitors significantly reduced the genomic methylation level and facilitated the formation of G4 structures.

**Figure 3.**
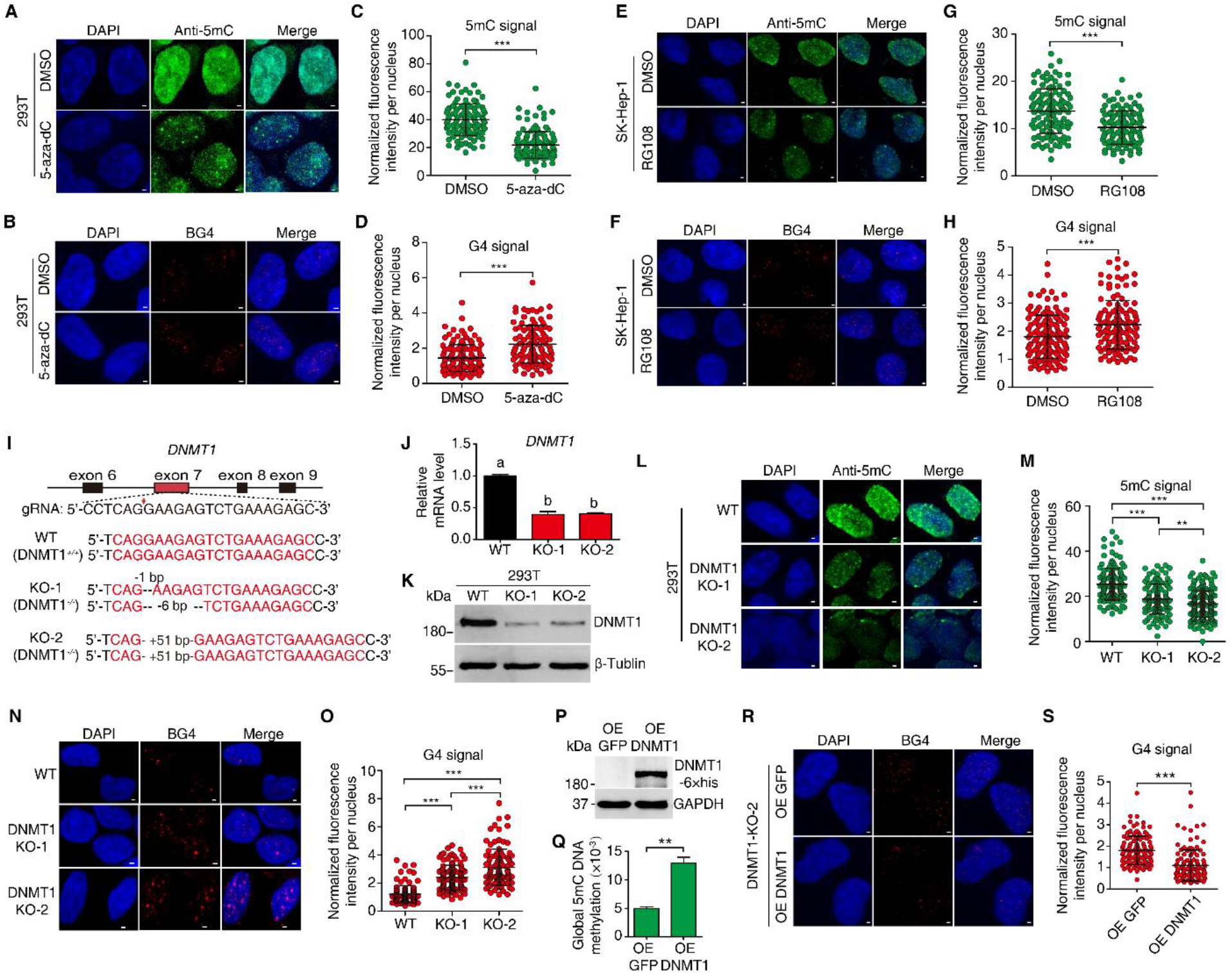
Effect of the reduced methylation level on the formation of the G4 structure. (A, E) Detection of methylation levels by immunofluorescence (IF) when cells were treated with 5-aza-dC and RG108. (B, F) The variation in G4 formation detected by IF using BG4 in the cells treated with 5-aza-dC and RG108. (C, D) Quantitative statistical analysis of the methylation level and G4 formation in the cells treated with 5-aza-dC. (G, H) Quantitative statistics of the methylation level and G4 formation in the cells treated with RG108. (I) Design of the knockout target sites and sequencing confirmation of the identified mutant strains. (J, K) Detection of the mRNA and protein levels of the DNMT1 gene in the wild type and mutants. (L-O) Detection and quantitative analysis of methylation levels (L and M) and the G4 signals (N and O) of the wild type and the mutants. (P) Detection of DNMT1 expression by western blot. (Q) The change in the methylation level when DNMT1 was compensatorily expressed in DNMT1-KO-2 cells. (R) Detection of the G4 signal when DNMT1 was compensatorily expressed in DNMT1-KO-2 cells. (S) Quantitative analysis of the immunofluorescence staining in R. Data are the mean ± SD (C, D, G, H, M, O and S) or mean ± SEM (J and Q). Statistical significance was determined by Student’s t test, **p<0.01, ***p<0.001. The scale bars equal 1 μm.

In mammals, DNA methylation is mediated by three major methylases, DNMT1, DNMT3a and DNMT3b. DNMT1 is critical for the maintenance of genome-wide methylation, while DNMT3a and DNMT3b are mainly responsible for *de novo* methylation (Jones 2012). Many studies have shown that the whole-genome methylation level can be significantly reduced when DNMT1 is knocked out (Li et al. 1992; Liao et al. 2015). In this study, DNMT1 knockout was performed in 293T cells by using CRISPR/Cas9 gene editing method, followed by detection of genomic methylation and G4 levels. Specific sgRNAs were designed to target exon 7 of the DNMT1 gene (Fig. 3I) and used to transfect 293T cells with a Cas9 plasmid to achieve DNMT1 knockout. Two mutant cell lines (DNMT1-KO-1 and DNMT1-KO-2) were obtained after monoclonal screening and confirmation by genomic DNA sequencing. In the DNMT1-KO-1 cell line, the DNMT1 alleles were mutated in an inconsistent manner. One allele had a 1 bp deletion, leading to an open reading frame shift mutation. The other allele lacked 6 bp nucleotides, resulting in two missing amino acid residues compared to the wild type (Fig. 3I). In the DNMT1-KO-2 cell line, both DNMT1 alleles gained 51 bp nucleotides at the same position, resulting in a 17-amino acid residue insertion (Fig. 3I). qPCR and western blot analysis proved that the expression of DNMT1 at both mRNA and protein levels significantly decreased in the two mutant cell lines compared to the wild type (Fig. 3J and K). The methylation level and G4 signal in the wild-type and two DNMT1 mutant cell lines were detected using an immunofluorescence assay, which showed that the overall methylation level was notably reduced in the two DNMT1 mutant cell lines, particularly in the DNMT1-KO-2 cell line (Fig. 3L and M), while the G4 signal was increased significantly in DNMT1 mutant cells, with more G4 signal in DNMT1-KO-2 cells (Fig. 3N and O). These results suggested that the decrease in the genomic DNA methylation level caused by the DNMT1 mutation led to the significant increase in G4 signal.

The above experiments demonstrated that DNMT1 knockout resulted in a decrease in the overall methylation level and an increase in the G4 formation. To further confirm this result, the effect of over-expressing DNMT1 in the DNMT1 knockout cell lines on the methylation level and G4 signal was examined. EGFP or DNMT1-EGFP was over-expressed in the DNMT1-KO-2 cells. The Western blot results showed that DNMT1 was over-expressed in the cells (Fig. 3P). The overall methylation level was significantly increased (Fig. 3Q), whereas the number of G4 structures was significantly reduced in the DNMT1-KO-2 cells over-expressing DNMT1 (Fig. 3R and S). These experiments demonstrated that G4 structure formation in the genome was strongly down-regulated by DNA methylation, but up-regulated by removal of methylation.

### DNA 5mC Modification Inhibited Folding of PQSs into G4s

To further assess whether DNA 5mC methylation affect G4 formation on a genome-wide scale, the abundance of G4 and 5mC in the genomes of wild type and DNMT1 knockout cells, DMSO and 5-aza-dC treated cells, was detected by DNA dot blotting assays. The results showed a reduction of the global genomic DNA 5mC methylation levels, with a global increase of G4 levels in the DNMT1 knockout cells (Fig. 4A and B) and in the 5-aza-dC treated cells (Fig. 4C and D).

**Figure 4.**
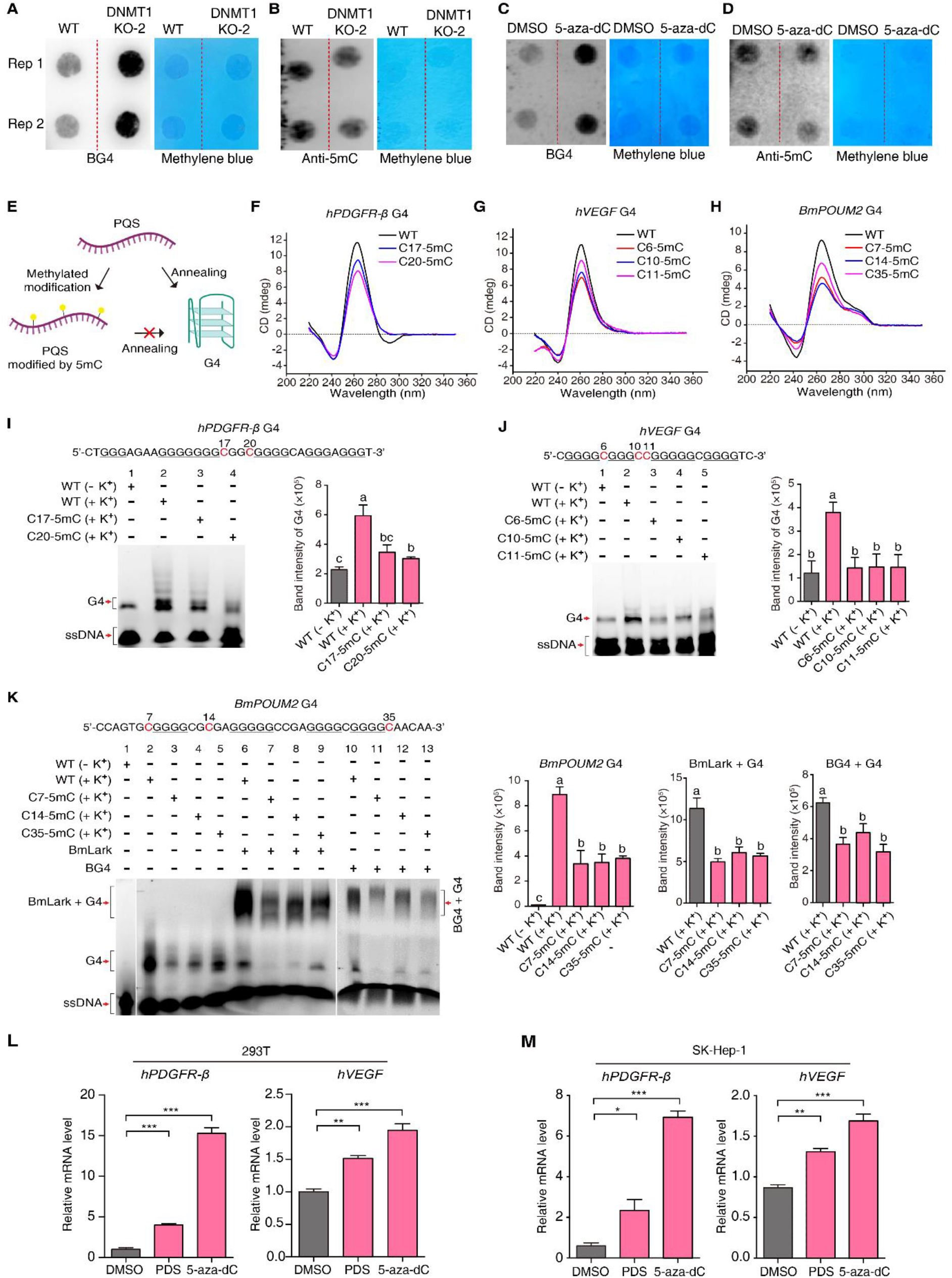
The effect of 5mC modification on G4 formation. (A-D) Dot blot assays for anti-5mC and BG4. The methylene blue staining of 200 ng total genomic DNA was used as loading control. (E) Descriptive diagram of the probe treatment method in CD and EMSA assays. (F-H) CD analysis of the effect of 5mC modification on G4 formation. (I, J) EMSA analysis and quantitative statistics of the effect of 5mC modification on G4 formation. (K) EMSA analysis and quantitative statistics of the effect of 5mC modification on G4 formation and its binding with protein. (L) qRT–PCR analysis of hPDGFR-β and hVEGF expression in 293T cells treated with PDS and 5-aza-dC. (M) qRT–PCR analysis of hPDGFR-β and hVEGF expression in SK-Hep-1 cells treated with PDS and 5-aza-dC. Data are the mean ± SEM. Statistical significance was determined by Student’s t test, *p<0.05, **p<0.01, ***p<0.001.

To demonstrate whether 5mC modification of PQSs directly inhibits G4 formation, three reported PQSs within the promoters of *hPDGFR-β* (Brown et al. 2017), *hVEGF* (Guo et al. 2008) and *BmPOUM2* (Niu et al. 2018, 2019) were selected to analyze the relationship between methylation and G4 formation. These PQSs have been proven to be capable of forming G4 structures either *in vitro* or *in vivo*. The methylation sites in the promoter PQS of *BmPOUM2* were tested by bisulfite sequencing PCR (BSP) analysis (Supplemental Fig. S1A). The methylation sites of the promoter PQSs of *hPDGFR-β* and *hVEGF* were obtained from whole-genome BSP data (Supplemental Fig. S1B and C). There are two 5mC modification sites located in the same loop of the *hPDGFR-β* PQS, three 5mC sites separated in two loops of the *hVEGF* PQS and three 5mC sites distributed in the 5’ end, first loop, and 3’ end of the *BmPOUM2* PQS (Supplemental Fig. S1). To investigate whether modification of these methylated site inhibits the G4 formation in these PQSs, PQS oligonucleotides with or without 5mC methylation were synthesized and used for G4 formation assay by the CD method (Fig. 4E). The results showed that the G4 feature absorption peaks of the 5mC-modified oligonucleotides were much lower than that of the unmodified oligonucleotides (Fig. 4F-H), indicating the inhibition of 5mC methylation on G4 formation as shown in Fig. 4E. This inhibitory effect was further confirmed by EMSA. In the presence of K^+^, more G4 structures formed in the *hPDGFR-β, hVEGF* and *BmPOUM2* PQS oligonucleotides than that in the absence of K^+^ (Fig. 4I-K lanes 1 and 2). The G4 bands notably decreased in intensity or even disappeared when these PQS oligonucleotides were methylated (Fig. 4I lanes 3 and 4, J lanes 3-5 and K lanes 3-5), suggesting that the 5mC methylation in the PQSs suppressed the formation of G4 structures. Furthermore, to verify the inhibitory effect of 5mC methylation on G4 folding and binding with proteins, the DNA-protein binding between the G4 and G4-binding proteins LARK and BG4 was examined. The results revealed that the intensity of the LARK-G4 (Fig. 4K lane 7-9) or BG4-G4 (Fig. 4K line 11-13) binding bands was reduced compared with that of the control (Fig. 4K lane 6 and 10), further proving that the 5mC-methylated PQSs resulted in fewer G4 structures.

It has been demonstrated that the *hPDGFR-β* and *hVEGF* promoters contain G4 structures (Guo et al. 2008; Brown et al. 2017). This study proved that the 5mC methylation could influence the formation of G4 structures in the promoter PQSs. Thus, it is logical to propose that the transcription of *hPDGFR-β* and *hVEGF* would be affected by the G4 stabilizer pyridostatin (PDS) or the methylation inhibitor 5-aza-dC. We treated 293T and SK-Hep-1 cells with PDS or 5-aza-dC, and then detected the expression of *hPDGFR-β* and *hVEGF*. Compared to the control treatment, both compounds significantly up-regulated the expression of the *hPDGFR-β* and *hVEGF* genes (Fig. 4L and M). This result indicated that either the stabilization of the G4 structure by PDS or the induction of G4 formation through 5-aza-dC-mediated methylation inhibition consistently activated the expression of *hPDGFR-β* and *hVEGF*. The upregulation effect of 5-aza-dC-mediated methylation inhibition on the transcription of the genes appeared more significant than that of the PDS treatment, suggesting that extra 5mC modification sites probably existed in the promoters of these genes in addition to the 5mC modification sites within the G4 sequences. Overall, both the G4 structures and 5mC methylation antagonistically regulated the transcription of these genes, with G4 structures facilitating and 5mC methylation suppressing their expression.

### Genomic Hypomethylation in Favour of Gene Transcription by Accommodating More G4s

In order to further verify the effects of methylation on G4 formation in human genome, we performed G4 CUT&Tag with wild type (WT) and DNMT1-KO-2 293T cells using BG4 antibody. We identified 16041 G4 peaks in WT and 14337 G4 peaks in DNMT1 KO-2 cells, respectively. The Venn diagram distribution revealed that 12322 peaks were shared in both WT and DNMT1 KO-2 (Fig. 5A). We next annotated all peaks and found that the majority of G4s localized at the gene promoter regions and higher percentage of G4 peaks located at the gene promoter regions in DNMT1 KO-2 cells (75.3%) than in WT cells (71.3%) (Fig. 5A). Moreover, the motif discovery showed that the top two statistically significant motifs were very similar in both DNMT1 KO-2 cells and WT cells (Fig. 5B and C). The enrichment of whole genome BG4 signal was significantly higher in DNMT1 KO-2 cells than in WT cells (Fig. 5D). Furthermore, the intensity of the BG4 signal within the 12322 common G4 peaks was also higher in DNMT1 KO-2 cells than in WT cells (Fig. 5E), suggesting that more G4 structures were present in hypomethylated cells.

**Figure 5.**
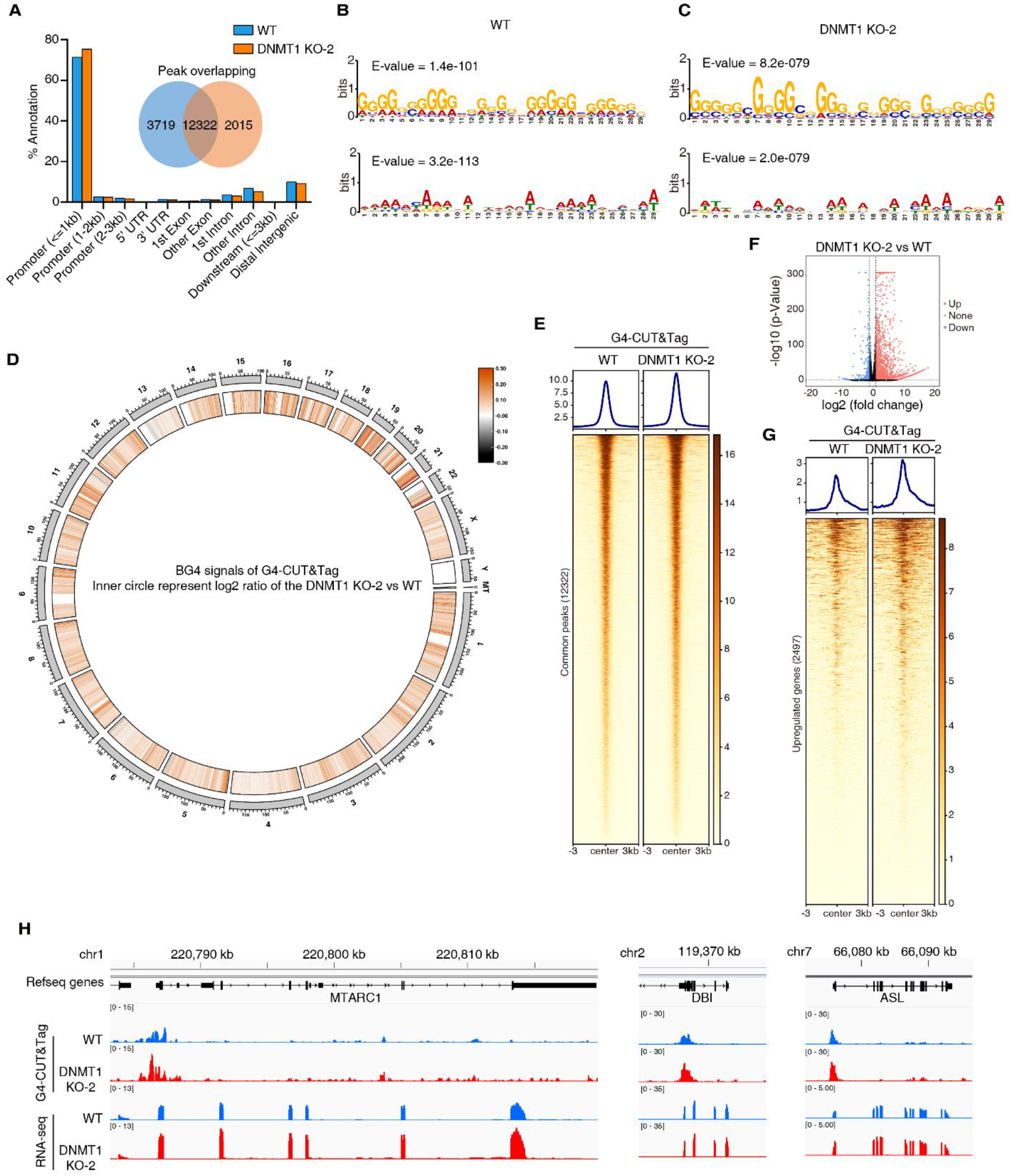
Hypomethylation enhances gene transcription by accommodating more G4 formation. (A) Genome-wide annotation of 293T WT and DNMT1 KO-2 G4-CUT&Tag peaks, presented as a percentage. Venn diagram shows the number of WT or DNMT1 KO-2 specific peaks and overlapping peaks. (B, C) MEME-chip analysis on the BG4 peaks of WT or DNMT1 KO-2 showing the top two motifs. (D) Circus plot showing genome-wide BG4 density. The outermost circle represents chromosomes, and inner circle represent log2 ratio of the DNMT1 KO-2 vs WT. In DNMT1 KO-2 and WT BG4 signals, IgG signals were used as the background for the deduction respectively and normalized by BPM (Bins Per Million mapped reads). (E) Heat map of G4-CUT&Tag enrichment from WT or DNMT1 KO-2 centered on common peaks (12322) with a ±3 kb. (F) Volcano plot for the comparison of RNA-seq data between WT and DNMT1 KO-2. No changed genes are shown in black, while differently expressed genes (fold-change > 2 and p-Value < 0.05) are denoted in blue or red. (G) Heat map of G4-CUT&Tag enrichment on DNMT1 KO-2 VS WT up-regulated gene (2497) promoter regions (TSS ±3 kb). (H) Genome browser tracks showing the distribution of G4 and RNA-Seq for WT (blue tracks) and DNMT1 KO-2 (red tracks). The black track and arrows at the top show the gene structures (5’UTR, exons, introns, 3’UTR) and the direction of transcription.

It has been known that G4s in promoter regions are associated with transcriptional activation (Halder et al. 2010; Hänsel-Hertsch et al. 2016) and hypomethylation in mammals (Yin et al. 2017). Therefore, we analyzed whether hypomethylation in human genome affects gene expression and its correlation with G4 formation. The RNA-seq with WT and DNMT1 KO-2 cells showed that when DNMT1 was knocked out, 3647 differentially expressed genes (DEGs) were identified, among which 2645 were up-regulated and 1002 were down-regulated (Fig. 5F). We statistically analyzed the BG4 signal in the promoter region of the 2497 up-regulated DEGs that have well-defined TSS, and found that the BG4 signal near the TSS (± 3kb) of the up-regulated DEGs increased in the cells with DNMT1 deletion (Fig. 5G). As examples, the MTARC1, DBI and ASL genes had a higher G4 enrichment in their promoters and a higher RNA-seq reads in DNMT1-KO cells (Fig. 5H). Taken together, these results demonstrated that the reduction of global genome methylation leads to the formation of more G4s, which elevates gene transcription levels.

## DISCUSSION

More than 700,000 PQSs were predicted in human genome by a high-throughput sequencing method (Chambers et al. 2015). Nevertheless, the number of detected G4 (dG4) by G4 ChIP-seq (Hänsel-Hertsch et al. 2016) or G4 CUT&Tag (Lyu et al. 2022) assay was much less than the number of predicted PQSs, suggesting that not all PQSs can form G4 structures *in vivo* and that specific cellular and physiological conditions are required for G4 formation. In human, over 40% of genes were predicted to contain one or more PQSs in their promoter regions using bioinformatic method (Huppert and Balasubramanian 2007), and the *bona fide* G4 structures intracellularly identified using BG4-ChIP-seq are prevalent in the promoter of active transcribed genes (Hänsel-Hertsch et al. 2016, 2020). All these studies suggest that PQSs can fold into G4 structures when the double-stranded DNA unwinds to form a single strand, for example, prior to gene transcription, although G4 formation was not necessarily coupled with transcription (Shen et al. 2021). However, the intrinsic mechanism of global G4 formation in a genome remains elusive.

Interestingly, an antagonistic relationship between G4 and methylation was found in our previous study (Wu et al. 2021), in which a mammalian species (pig) with a highly methylated genome and an insect species (silkworm) with relatively low methylation were used to examine the relationship between G4 and methylation in the upstream 2 kb regions of genes. The results showed an antagonistic relationship between G4 and methylation, which is conserved in evolution (Wu et al. 2021). It inspired us to further confirm this negative correlation and whether DNA methylation can affect the formation of G4 structures. High-throughput identification of G4 structures based on BG4- and G4-specific binding probes (G4P) has been established, including BG4-ChIP-seq (Hänsel-Hertsch et al. 2016), BG4-CUT&Tag (Lyu et al. 2022) and G4P-seq (Zheng et al. 2020), by which a large amount of G4 identification and distribution data was obtained, making it possible to draw a G4 landscape at the whole-genome level. Combining the WGBS and BG4-CUT&Tag analyses, a strong and negative correlation between dG4 and methylation of genomic DNA was found in this study (Fig. 1). Even in the condition of the similar contents of PQSs, hypermethylated sequences formed fewer G4 structures than hypomethylated sequences. Furthermore, the same PQS sites had fewer G4 structures when hypermethylated but had more G4 structures when hypomethylated (Fig. 1G and H). Similar negative correlation was also found in cancer tissues and cell lines. Such a wide and conserved existence of this negative relationship between G4 formation and DNA methylation makes it more intriguing to investigate the intrinsic mechanism. This phenomenon can be explained by two possible scenarios: the G4 structure prevents DNA methylation, or DNA methylation inhibits G4 formation. Inhibition of the G4 structure from DNA methylation has been reported in several studies (Halder et al. 2010; Mao et al. 2018). However, there is no report on whether DNA methylation inhibits G4 formation. This study provides experimental evidence to demonstrate the inhibitory effect of DNA methylation on G4 formation. When the global genomic methylation level was reduced by methylation inhibitor treatment or DNMT1 knockout, the overall genomic G4 signals were increased (Fig. 3). When the expression of DNMT1 was rescued by complementarily expressing DNMT1 in the DNMT1 knockout cells, the G4 structures were correspondingly decreased (Fig. 3). Furthermore, the PQS oligonucleotides that were proven to be able to fold into G4 structures in the presence of K^+^ could hardly form G4 structures when they were 5mC methylated (Fig. 4). Methylation inhibitor treatment or G4 stabilizer treatment affected gene expressions consistently (Fig. 4). These results strongly suggest the inhibitory effect of DNA methylation on G4 formation.

In most cases, methylation inhibits gene transcription by several mechanisms; for example, methylated DNA inhibits its binding with transcription factors; methylated DNA recruits CpG-binding proteins to form transcription-inhibiting complexes; or methylation changes the chromatin architecture and affects gene transcription. In this study, increased methylation inhibits the formation of G4, turning off gene transcription, whereas decreased methylation promotes the formation of G4, turning on gene transcription. This provides a novel mechanism to the general inhibitory effect of DNA methylation on global genomic transcription.

Similar to DNA methylation, G4 is considered a novel epigenetic regulatory mechanism for gene transcription and phenotype formation because both DNA methylation and G4 formation are independent of first nucleotide sequences. This study explored the relationship between these two epigenetic mechanisms, particularly the effect of DNA methylation on G4 formation in genomes, and clearly demonstrated that they are antagonistic to each other. It is worth to notice that the interaction between DNA methylation and G4 formation occurs at DNA molecules, rather than at chromatin. In the *in vitro* experiments such as dot blot, circular dichroism and EMSA, in which the purified genome DNA or synthesized DNA oligonucleotides without chromatin protein were used. In dot blot assay, when the methylation levels were reduced by inhibitors or DNMT1 knockout, the G4 structures increased (Fig. 4A-D). In circular dichroism (Fig. 4F-H) and EMSA (Fig. 4I-K) assays, the 5mC modified PQSs form much less G4 structures. All these *in vitro* assays demonstrate the direct effect of DNA 5mC methylation on G4 formation, not through the chromatin structure. This antagonistic relationship can be explained by the features of the chemical structures of DNA methylation and G4 structure. Firstly, both methylation and G4 structures usually occur in the GC-rich regions of genomes. Methyl groups added to cytosines within a PQS by methylase may form a steric hindrance impeding G4 formation. Secondly, DNA methylation may affect the chromatin accessibility and further influences DNA G4 formation. Thirdly, methylated DNA may block nuclear proteins that are required for G4 formation to bind with PQS, or lose the capability of binding with such proteins. Conversely, the folded G4 structure may block the interaction of methyltransferase with DNA so that methylation could not take place. All these factors could contribute to the antagonistic relationship between these two epigenetic regulation mechanisms of gene transcription. The detail chemical and structural mechanism underlining this antagonistic relationship remains to be further investigated.

Many studies have shown that G4 structure formation is frequently encountered in human diseases, especially oncogenesis (Balasubramanian et al. 2011; Neidle 2016; Nakanishi and Seimiya 2020). G4 abundance is significantly increased in gastric cancer and liver cancer tissues (Biffi et al. 2014) and G4 abundance and location have been used to classify 22 breast cancer patient-derived tumor xenograft (PDTX) into at least three G4-based subtypes (Hänsel-Hertsch et al. 2020). However, the molecular mechanism of the increased G4 abundance in tumor tissues remains to be expounded. In this study, similar G4 pattern was found in the tested cancer cells. G4 abundance was significantly increased in colon cancer tissues and liver cancer cell line, meanwhile the global methylation level was reduced (Fig. 2). It was consistent with the reports, in which global genomic methylation was reduced in stomach cancer (Guo and Yan 2015) and liver cancer (Guerrero-Preston et al. 2007). Consequently, we speculate that decreased DNA methylation in tumor tissue contributes the increase in G4 abundance, leading to the unprogrammed expression of oncogenes, and finally to oncogenesis. Thus, if it is true, G4 structures can be considered as a possible molecular target for tumor diagnosis and therapy.

A number of studies have showed that the G4 structures facilitate to gene transcription (Hänsel-Hertsch et al. 2016; Li et al. 2021), whereas DNA methylation generally inhibits gene transcription (Smith and Meissner 2013). Here, we propose a mechanism model (Fig. 6) based on our results. Transcription of the genes with PQSs in their promoter regions is inhibited when the PQS sites are 5mC methylated, which inhibits the formation of G4. In certain circumstance, such as in cancer tissues, the global methylation rate is decreased, resulting in more G4 formation, which in turn elevates gene transcription. This model suggests a regulatory mechanism of gene transcription through the interaction between G4 structure and methylation of genomic DNA.

**Figure 6.**
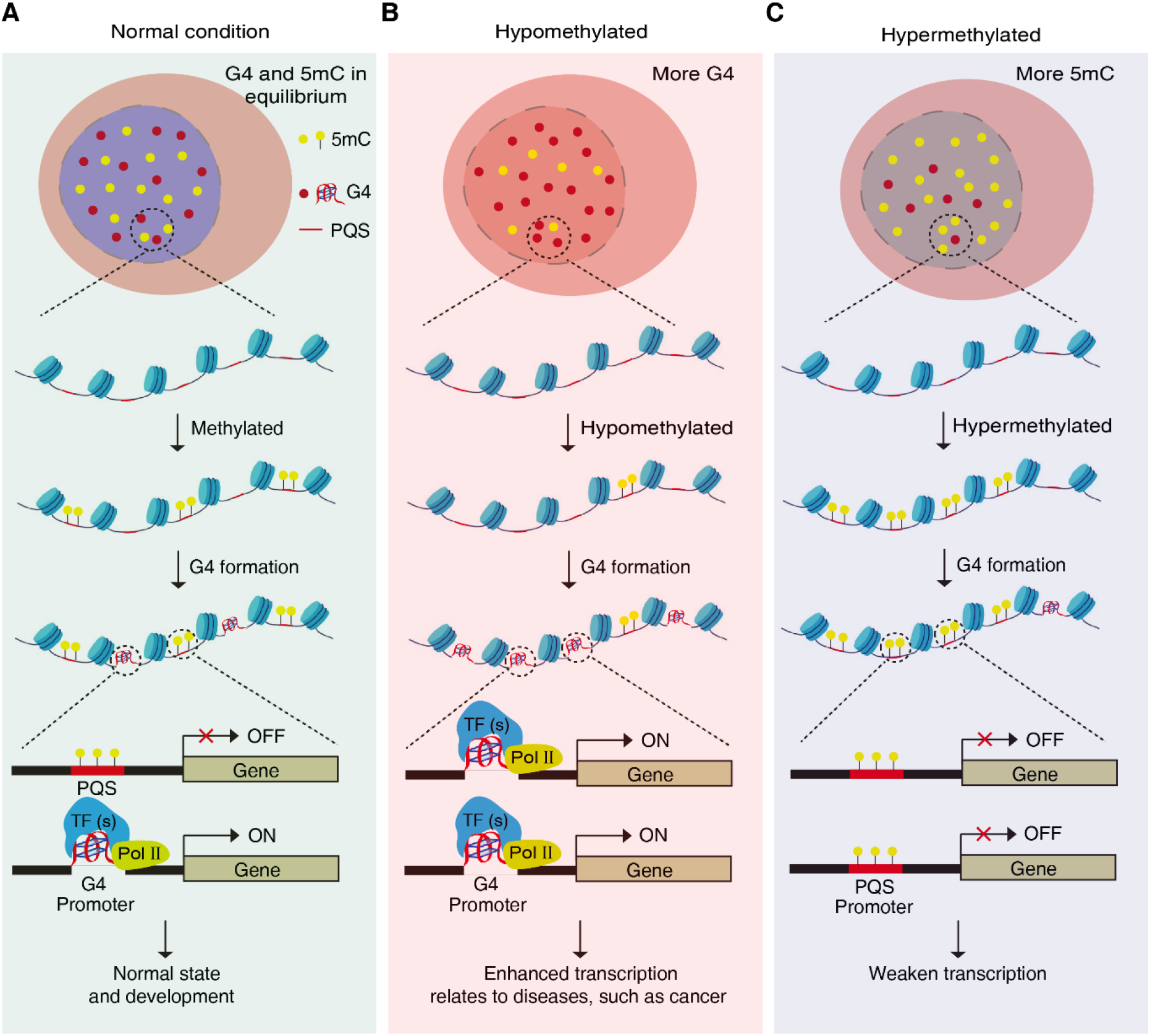
Diagram of proposed epigenetic regulatory mechanism of DNA G4 structures and 5mC methylation in gene transcription. In normal condition, the global gene transcription is in equilibrium. Genes with PQSs that are not methylated are active transcribed due to the formation of G4, while the transcription of genes with methylated PQSs are inhibited (A). In some cases, such as cancer, the overall genome is hypomethylated and more G4 structures are formed which resulted in elevated transcription, especially oncogenes (B). However, when cell genome is hypermethylated, the G4 formation is inhibited and meanwhile the overall gene transcription in weakened (C).

## METHODS

### Cell Culture, Chemical Treatment and Clinical Samples

Human 293T cells (CRL-3216), a derivative of human embryonic kidney 293 cells, SK-Hep-1 human hepatic adenocarcinoma cells (HTB-52) and HepG2 cells (HB-8065) were purchased from ATCC (American Type Culture Collection, VA, USA) and cultured according to the instructions. All cell lines were maintained in DMEM cell medium (11995065, Thermo Scientific, MA, USA) containing fetal bovine serum (FBS, 10099141, Thermo Scientific, MA, USA), penicillin and streptomycin (15140122, Thermo Scientific, MA, USA) with a final concentration of 10%, 100 U/mL and 100 μg/mL respectively. Cells were incubated in a 37 °C incubator with 5% CO_2_. Mycoplasma was tested periodically to eliminate mycoplasma contamination with all used cells. The compounds 5-aza-dC (5-Aza-2’-deoxycytidine, A3656, Sigma, Shanghai, China) and RG108 (N-phthalyl-L-tryptophan, HY-13642, MedChemExpress, NJ, USA) were dissolved in DMSO (dimethyl sulfoxide). The 293T and SK-Hep-1 cells grown on glass coverslips were treated with 5 μM 5-aza-dC or 150 μM RG108 in duplicate cultures with the supplemented medium changed every 24 h over three days. The clinical tissue samples were collected from Guangzhou Panyu Central Hospital.

### Immunofluorescence Staining

Immunofluorescence assays of G4s were conducted as described in previous studies (Hänsel-Hertsch et al. 2016). Cells were seeded on glass coverslips and cultured overnight at 37 °C. Cell fixation was performed by incubating cells with 4% paraformaldehyde for 10 min at room temperature (RT). Then cells were permeabilized with 0.2% Triton X-100 in PBS for 10 min at RT. After twice rinse with PBS, cells were blocked for 1 h at 37 °C in a blocking solution composed of 5% bovine serum albumin (BSA) and 5% goat serum in phosphate-buffered saline (PBS). After blocking, the cells were incubated with a BG4-Flag antibody (1:100 diluted in 0.5% goat serum + PBST, MABE917, Merck, Darmstadt, Germany) for 1 h at 37 °C, followed by three washes with PBST for 5 min each. The cells were incubated with a 1:800 dilution of rabbit anti-Flag antibody (2368, Cell Signaling Technology, MA, USA) for 1 h at 37 °C, followed by washing three times with PBST for 5 min each. The cells were then incubated with 1:500 Alexa Fluor 594 secondary antibody (A11037, Thermo Fisher, MA, USA) for 45 min at RT, followed by washing three times with PBST for 5 min each and a second wash containing DAPI (2.5 μg/mL). Coverslips were mounted on the glass slide with ProLong Diamond anti-fade mounting medium (P36970, Thermo Fisher, MA, USA). The immunofluorescence assay of 5mC was conducted as described in a previous study (Santos et al. 2002). Similarly, cells grown on coverslips were fixed in 4% paraformaldehyde for 10 min and permeabilized with 0.2% Triton X-100 for 10 min at RT. Then, cells were treated with 2 M HCl for 30 min at RT and subsequently neutralized for 10 min with 100 mM Tris-HCl buffer, pH 8.5. The cells were blocked in blocking solution (5% BSA and 5% goat serum in PBS) for 1 h at 37 °C. After blocking, the cells were incubated with anti-5mC antibody (1:200 diluted in 0.5% goat serum + PBST, 39649, Active Motif, CA, USA) at 4 °C overnight and then washed three times with PBST for 5 min each. The cells were then incubated with a 1:500 dilution of Alexa Fluor 488 secondary antibody (A11029, Thermo Fisher, MA, USA) for 45 min at RT and then washed three times with PBST for 5 min each, with the second wash containing DAPI (2.5 μg/mL). Finally, coverslips were placed on glass slides with ProLong Diamond anti-fade mounting medium (P36970, Thermo Fisher, MA, USA). Immunofluorescence image acquisition was conducted using Olympus Fluoview FV1000 confocal microscope. Image Z stacks were obtained using the ‘pinhole’ set to 0.6 Airy units. The same parameters were applied to all the compared samples.

Quantitative statistical analysis of confocal images was performed using ImageJ software. The signals of 100-200 nuclei from three replicates were counted for each condition. Data are shown as the mean ± standard deviation (SD). Statistical significance between the samples was determined by Student’s *t test* using GraphPad Prism version 5. The fluorescence intensity values were normalized to the nuclear area.

### Knockout of DNMT1 in Cells

To establish knockout cell lines, the genome editing one vector system (PX459) (104142, Addgene, MA, USA) was used. The sgRNAs targeting *DNMT1* were designed using the CRISPR tool (http://crispr.mit.edu). To construct the plasmid PX459-gRNA, a short single-strand DNA containing a partial reverse complement of the gRNA sequence was synthesized. After annealing, the DNMT1 gRNA with a sticky end containing the *Bbs* I site was formed. The gRNA sequence was annealed to the plasmid PX459 and digested with *Bbs* I, generating the recombinant plasmid PX459-DNMT1-gRNA. The PX459-DNMT1-gRNA plasmid was then used to transfect 293T cells using FuGENE HD Transfection Reagent (E2311, Promega, WI, USA) according to the manufacturer’s instructions. An equal amount of PX459 plasmid DNA was used as a negative control. At 48-72 h after cell transfection, puromycin medium was used to screen knockout-positive cells. The knockout-positive cell populations were inoculated into 96-well plates by limiting dilution analysis. When the cells gradually grew from a 96-well plate to a 6-well plate, genomic DNA was extracted from the cells, and PCR was performed, followed by TA cloning. At least five clones were selected for sequencing verification until homozygous knockout cell lines were obtained for protein expression and functional analysis.

### Quantitative Real-Time PCR (qRT–PCR) and Western Blotting (WB)

qRT–PCR and WB were performed as described in our previous studies (Niu et al. 2019). Total RNA was extracted from 293T cells using an Eastep Super Total RNA Extraction Kit (LS1040, Promega, WI, USA). Reverse transcription was conducted using GoScriptTM Reverse Transcription Mix and Random Primers (A2800, Promega, WI, USA) according to the manual. The whole cell lysate were obtained with RIPA and the protein concentration was measured using BCA protein assay kit (23225, Thermo Scientific, MA, USA). Monoclonal antibody anti-DNMT1 (ab188453, Abcam, Cambridge, UK) was used to detect DNMT1 with a dilution of 1:1000. Polyclonal antibodies anti-His (ab9108, Abcam, Cambridge, UK) and anti-GAPDH (ab9485, Abcam, Cambridge, UK) were used to detect DNMT1-His and GAPDH at a dilution of 1:1000 and 1:3000 respectively.

### Cell Transfection

293T-KO-2 cells were inoculated in 12-well culture plates at desired density and cultured in DMEM (11995065, Thermo Fisher, MA, USA) overnight. Cell transfection was performed with Fugene HD transfection reagent (E2311, Promega, WI, USA) as described previously (Niu et al. 2019). Briefly, when the cell density reached 80%, a mixture of 100 μL which contained 2 μg of pEGFP (control) or pEGFP-DNMT1, 6 μL Fugene HD transfection reagent and appropriate volume of Opti-MEM Reduced Serum Medium (31985070, Thermo Fisher, MA, USA) was prepared and incubated for 15 min at RT. Then, the mixture was added to the 293T-KO-2 cells in DMEM and cultured for 48 h at 37 °C before G4 structure immunostaining.

### Frozen Sections

The specimens were embedded in freezing compound (Tissue-Tek O.C.T. Compound; Sakura Finetek-USA, USA) and fixed in a precooled specimen disc. Each specimen was frozen at - 20 °C using a Leica CM1950 cryostat (Leica CM1950; Leica Biosystems Nussloch GmbH, Germany) and trimmed until the specimen appeared on the cutting surface. Specimens were cut into 5-7 μm sections.

### Circular Dichroism (CD)

The wildtype and 5mC modified oligonucleotides were synthesized by Qingke Biotechnology (Beijing, China). The oligonucleotide sequences are shown in Supplemental Table S1. Oligonucleotides at 5 μM in 50 mM Tris-HCL (pH 7.5) and 100 mM KCL were heated at 95 °C for 10 min and cooled slowly to RT. CD experiments were performed as described in our previous study (Niu et al. 2019).

### Electrophoretic Mobility Shift Assay (EMSA)

The EMSA experiments were performed as described in our previous study (Xiang et al. 2022). The wildtype and 5mC modified oligonucleotides were labeled with 6-FAM at 5’ terminus by Qingke Biotechnology and annealed to form G4 probes as described previously (Xiang et al. 2022). The EMSA assay was conducted with 2 μM probes. In the binding reaction, 2 μg purified recombinant LARK protein or 1 μg BG4 (MABE917, Merck, Darmstadt, Germany) was added. The sequence of oligonucleotide probes used in this study are shown in Supplemental Table S1.

### Bisulfite Sequencing PCR (BSP) Analysis

*B. mori* genomic DNA was extracted from 2-day-old 2^nd^ instar larvae. Unmethylated cytosines were converted into uracil by using a MethylDetector kit (55001, Active Motif, CA, USA), whereas methylated cytosines remained unchanged. PCR was then performed with primers designed based on the BmPOUM2 G4 sequence. The PCR products were sequenced to confirm the 5mC modified sites.

### Dot Blotting Assays

For the dot blotting assay, genomic DNA was extracted from WT or mutant, DMSO or 5-aza-dC-treated cells (51304, Qiagen, DUS, GER). 200 ng DNA was dropped onto a positively charged Nylon membrane (Amersham Biosciences, Boston, USA). For G4 assays, membranes were allowed to air dry and then blocked in 5% milk for 1 h at RT. Then the membranes were incubated with BG4 (1:500) overnight at 4 °C followed by incubation with anti-FLAG antibody (1:1000) for an additional 1 h at RT and the secondary antibody (1:5000) for 1 h at RT, and ECL was applied and film was developed. For 5mC assays, the samples were heated at 95 °C for 10min to denature DNA. Then the membranes were incubated with anti-5mC (1:1000) overnight at 4 °C followed by incubation with the secondary antibody (1:5000) for 1 h at RT, and Immunoreactivity was detected using enhanced chemiluminescence (ECL) (32209, Thermo Fisher, MA, USA). The membranes were washed with TBST three times, 5 min each, before each procedure. After imaging, the membranes were stained with 0.02% methylene blue as loading control.

### Cleavage Under Targets and Tagmentation (CUT&Tag)

CUT&Tag assay was performed with Hyperactive Universal CUT&Tag Assay Kit (Vazyme, Nanjing, China) as described in our previous study (Xiang et al. 2022). Wildtype and DNMT1 KO-2 293T cells (1 × 10^6^) were used as materials and 5 μg BG4 or IgG (contorl) was used in the CUT&Tag assay. Data analysis was conducted as described in Li et al. (2021). In the motif analyses, only the top two significantly enriched motifs with the highest E-values were listed in the text.

### RNA-seq Processing and Analysis

Each sample was repeated three times. We confirmed the quality using FastQC and processed raw data by Trim Galore. We aligned the paired-end reads to human genome (UCSC-hg38) using Hisat2. A count table file indicating the number of reads per gene in each sample was generated using FeatureCounts. Differentially expressed genes were identified using DESeq and genes with a P-value <0.05 and a |log2FC| value >1 were considered to be differentially expressed.

### Bioinformatics Analysis

The location information of the human transcription start site (TSS) was downloaded from UCSC Genome Browser (http://genome.ucsc.edu). ComputMatrix was used to split into 10 bp windows to calculate the DNA methylation levels 1 kb upstream and downstream of the TSS (Ramírez et al. 2016). The average DNA methylation ratios of all cytosine sites in each window were used to represent the DNA methylation levels. The 10,000 transcripts with the highest methylation levels and 10,000 transcripts with the lowest methylation levels were selected, and the transcripts belonging to the same gene were eliminated to reduce redundancy. These genes were used to calculate the G4 signal strength within 1 kb both upstream and downstream of the TSS at 10 bp windows by using computeMatrix. Whole-genome bisulfite sequencing datasets for K562 (ENCSR765JPC) and HeLa (ENCSR550RTN) cells were downloaded from ENCODE (https://www.encodeproject.org/), and the BG4 CUT&Tag data for K562 and HeLa cells were obtained from GSE178668 (Li et al. 2021).

## DATA ACCESS

Raw data for RNA-seq and CUT&Tag are available in the GEO repository under the accession number GSE221437. All data reported in this paper will be shared by the lead contact upon request. Original western blot (Supplemental Fig. S2 and S3) and EMSA images (Supplemental Fig. S4) have been added in Supplementary material.

## ACKNOWLEDGMENTS

This work was supported by National Natural Science Foundation of China (31930102; 32000337; 31720103916; 32102610) and China National Postdoctoral Program for Innovative Talents (BX20190123). We would like to thank BioRender (https://biorender.com/) for the creation of diagram.

## AUTHOR CONTRIBUTIONS

Q.F. and K.N. conceived and supervised the study. K.N., L.X. and X.L. performed experiments and related analyses. L.X. and H.X. collected genomic and methylomic data and generated bioinformatic analyses. Y.L., C.Z., J.L., J.L. X.Z., Y.P. and G.X. provided help in data collection. K.N., X.Z. and Q.F. wrote the manuscript. Q.F. and Q.S. review and editing the manuscript. H.W. provided tissue samples and conducted pathological analysis of samples. All authors edited and approved the final manuscript.

